# Patterns of population genomic diversity in the invasive Japanese knotweed species complex

**DOI:** 10.1101/2020.08.06.226886

**Authors:** Acer VanWallendael, Mariano Alvarez, Steven J. Franks

**Affiliations:** Fordham University, 441 E. Fordham Road, Bronx, NY 10458; Duke University, 130 Science Drive, Durham, NC 27708

**Keywords:** invasive species, genetic diversity, Japanese knotweed, clonal, polyploid

## Abstract

**Premise:** Invasive species are expected to experience a reduction in genetic diversity due to founder effects, which should limit their ability to adapt to new habitats. Still, many invasive species achieve widespread distributions and dense populations. This paradox of invasions could potentially be overcome through multiple introductions or hybridization, both of which increase genetic diversity. We conducted a population genomics study of Japanese knotweed (*Reynoutria japonica*), which is a polyploid, clonally reproducing invasive species that has been notoriously successful worldwide despite supposedly low genetic diversity.

**Methods:** We used Genotyping-by-Sequencing to collect 12,912 SNP markers from 88 samples collected at 38 locations across North America for the species complex. We used non-alignment based k-mer hashing analysis in addition to traditional population genetic analyses to account for the challenges of genotyping polyploids.

**Results:** Genotypes conformed to three genetic clusters, likely representing Japanese knotweed, Giant knotweed, and hybrid Bohemian knotweed. We found that, contrary to previous findings, the Japanese knotweed cluster had substantial genetic diversity, though it had no apparent genetic structure across the landscape. In contrast, Giant knotweed and hybrids showed distinct population groups. We did not find evidence of Isolation-by-Distance in the species complex, likely reflecting the stochastic introduction history of this species complex. Among species, we found no correlations between SNPs and several temperature- and precipitation-based climatic variables.

**Conclusions:** The results indicate that clonal invasive species can show substantial genetic diversity and can be successful at colonizing a variety of habitats without showing evidence of local adaptation or genetic structure.

## Introduction

Understanding genetic changes accompanying population expansion is critical for predicting ecological and evolutionary dynamics in colonizing species. Organisms colonizing a new area often experience a type of population bottleneck known as a founder effect, in which a subset of individuals starts a new population with reduced genetic diversity compared to the original population (Mayr 1942). As the population expands in the new regions, serial founding events may repeat across the landscape (Klopfstein et al. 2005; Excoffier & Ray 2008), magnifying the effect. Resultant genetic drift during colonization may lead to reduced genetic diversity (Wright 1931; Dlugosch and Parker 2008; Excoffier and Ray 2008), which may in turn lead to loss of beneficial traits as well as a reduction in the efficacy of selection (Frankham 1995; Peischl et al. 2013; Blackburn et al. 2015). Invasive species, typically defined as non-native organisms that spread rapidly and displace natives, represent a magnified example of population expansion. In what has been termed the “genetic paradox of invasion” (Frankham 1995; Allendorf & Lundquist 2003; Estoup et al. 2016), many introduced species, despite the predicted negative effects of the loss of genetic diversity, are highly successful and become invasive. Invasive plants provide a substantial threat to biological diversity, and their ecology and evolution has been studied for decades (Butchart et al. 2010; Baker & Stebbins 1965), but researchers still know relatively little about the genomics of invasive plants, including the extent and effect of population genetic bottlenecks and the influence of factors such as clonal reproduction and hybridization on shaping responses to selection, and invasion success, across the landscape (Lee 2002, Chown et al. 2015, Allendorf and Lundquist 2003).

The perennial, dioecious plant *Reynoutria japonica* has been described as one of the “world’s worst invasive alien species” for its aggressive nature and challenges in its removal (Lowe et al. 2000). Since it is widespread in its introduced range, and reproduces through both sexual and asexual means, it represents a useful system to test hypotheses about the impacts of colonization on genetic diversity. The species is native to Japan and introduced to China, Australia, Europe, and North America. In North America, the first known introduction occurred in 1868 from a European source (Del Tredici 2017). Many subsequent introductions and explosive growth have spread the species in high densities around temperate North America (Barney 2006). Octoploid *R*. *japonica* forms a species complex with tetraploid *R*. *sachalinensis* (F. Schmidt ex Maxim.) Nakai (Giant knotweed) to form hybrid hexaploid *R*. x *bohemica* Chrtek & Chrtková (Bohemian knotweed; Bailey 2013). Both *R*. *sachalinensis* and hybrid *R*. x *bohemica* are sympatric with *R*. *japonica* in North America, although their density is difficult to estimate because many reports misidentify hybrids as *R*. *japonica* (Zika & Jacobson 2003; Gammon et al. 2007; Gaskin et al. 2014).

Notably, *R*. *japonica* has often been used as the quintessential example (Durka et al. 2005; Geng et al. 2007; Barrett et al. 2008; Herman & Sultan 2011) of an invasive species that is successful over a wide range of habitats and environments despite its supposed partial or complete lack of genetic diversity. *R*. *japonica* overwinters as a rhizome, and pieces of rhizome that fragment will grow into clones of the parent plant (Hollingsworth & Bailey 2000; Barney 2006; Richards et al. 2008). In the native range, there is substantial chloroplast haplotype diversity (Inamura 2000), but in the introduced range, *R*. *japonica* often has far less. For example, a study using RAPDs (Randomly Amplified Polymorphic DNA) indicated that all *R*. *japonica* in Great Britain is one female clone thought to be “one of the world’s largest vascular plants” (Hollingsworth & Bailey 2000). In North America, reports of genetic diversity have been mixed. There is strong evidence through chloroplast markers that there are several haplotypes, and that sexual reproduction is widespread (Grimsby et al. 2007; Grimsby & Kesseli 2010; Gammon et al. 2010, Gammon & Kesseli 2010). However, more recent AFLP (Amplified Fragment Length Polymorphism) studies have found genetic diversity only in *R*. *sachalinensis* and *R*. x *bohemica*, but none in *R*. *japonica* collected from both the Mid-Atlantic and Pacific Northwest regions (Richards et al. 2012; Gaskin et al. 2014). This difference in conclusions may be either due to a difference in sampling range or differences in the density and information content of the markers used. Therefore, we predicted that an increased number of markers and broader sequencing over a wide geographic range would help to clarify the distribution of genetic diversity present in North America.

The adaptive consequences of asexual reproduction are crucial to understanding the Japanese knotweed invasion. As colonizers, selfing and clonal plants possess several inherent advantages over obligate outcrossers, which could help explain why the capacity for uniparental reproduction is more common in invasive species than in those that do not become invasive (Pannell 2015; Razanajatovo & van Kleunen 2016). These advantages include a lack of need for other mates (Baker 1955), genetic continuity across generations that helps provide continued success in a given environment (Baker & Stebbins 1965), and reduced gene flow, which fosters local adaptation (Kawecki & Ebert 2004; Baker 1955; Baker & Stebbins 1965). But a disadvantage of asexual reproduction is that genetically limited populations that may be less able to adapt to novel biotic and abiotic stress (Crow & Kimura 1965; Verhoeven et al. 2010). It is well-established that *R*. *japonica* typically reproduces asexually, while *R*. *sachalinensis* and hybrids tend to spread through outcrossed seed (Bailey 2013). However, the population-level impacts of these reproductive systems on the distribution of genetic diversity have not yet been assessed.

This study examines the population and landscape genetics of the Japanese knotweed species complex in North America with the goal of understanding the factors that have permitted its high fitness despite founder effects. We used Genotyping-by-Sequencing (GBS) for 88 samples at 38 collection sites to capture 12,912 SNPs. Specifically, we sought to determine whether or not genetic diversity has been limited in Japanese knotweed due to founder effects combined with clonal propagation. Further, we hypothesized that genetic diversity would be partitioned differentially within the species complex, with higher genetic diversity in *R*. *sachalinensis* and hybrids. Finally, we examined the geographic dispersal of genotypes and their correlation with climatic variables to test whether founder effects have limited the adaptive potential of knotweed in the invaded range. The results of this work will provide insight into population genetic processes of species that can successfully colonize a wide variety of habitats.

## Materials and Methods

### Study design

We structured our sampling to examine diversity at multiple geographic scales to understand the landscape and microgeographic structure of populations. We did not make *a-priori* distinctions between hybrid species in the field to avoid biased sampling. We collected fresh leaf or rhizome tissue from 38 sites in the eastern portion of the United States (Figure 1; Table S2). When possible, fresh leaves were collected and frozen within one hour at −20^°^C until extraction. At more distant sites, we collected rhizomes into a cooler, then germinated them in the greenhouse to obtain fresh leaves. Within sites, we collected several individuals and sequenced at least two individuals per site whenever possible. As *R*. *japonica* spreads mainly via rhizomes, groups of “individuals” may in fact be one organism. Since distinguishing individuals is impossible barring excavation of the rhizome network, we followed the suggestions of Richards et al. (2008) and collected only from rhizomes spaced 10 m apart. For Midwest and West Coast samples, leaves were collected and dried by citizen scientists then sent by mail. During collection, we assigned each stand to one of four habitat types: riparian, roadside, forest edge, or lacustrine. Riparian, lacustrine, and roadside habitats were all within 50 m of a river, lake, or road, respectively. All other sites were found at the border (forest edge) between a wooded area and an anthropogenic landscape. We recorded the sex of each individual based on floral characteristics and used the protocol of Zika & Jacobson (2003) to assign plants to a species.

**Figure 1.**
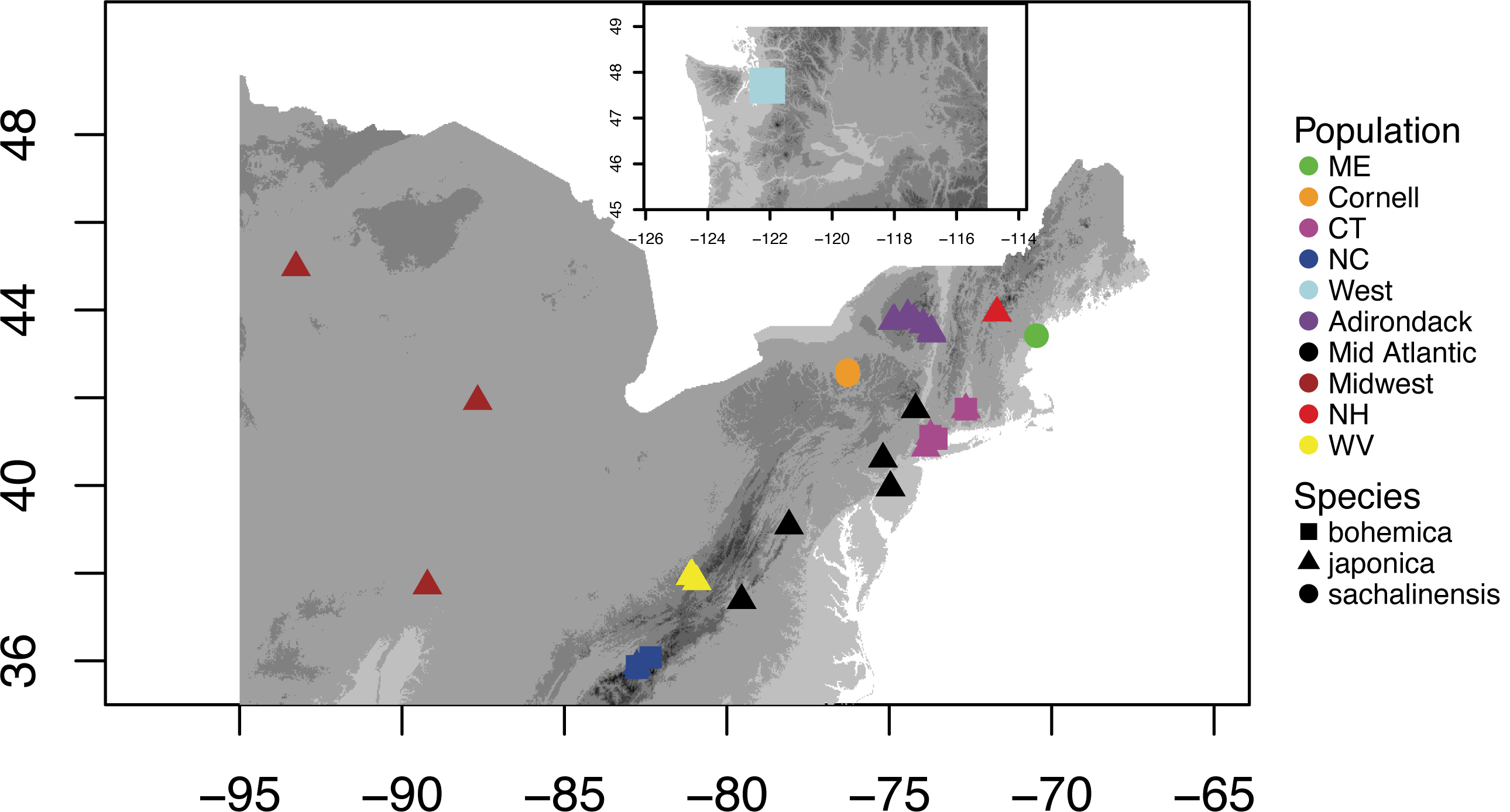
Sample collection locations in eastern North America. Population identity is shown by color, species is shown by shape. Sites CT and NC contained both *R*. x *bohemica* and *R*. *sachalinensis*. Groups used in analyses are shown by color. Inset shows the single West coast sample, collected from Seattle, WA.

**Figure 2.**
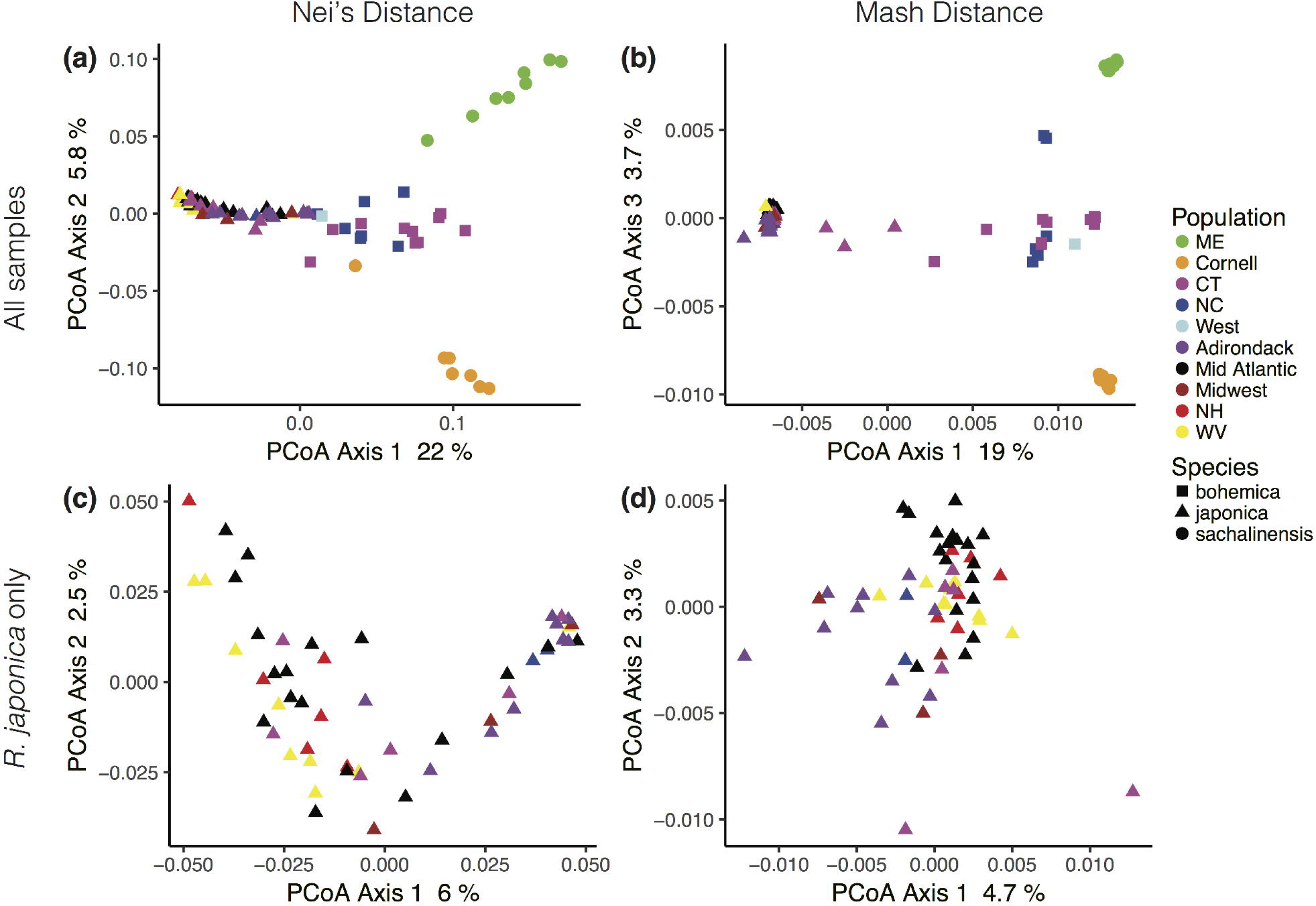
Principal Coordinates Analysis of distance matrices produced by two methods. Projected inertia is shown with the axis labels. Species identification was based on morphological differences. (a) Nei’s distance matrix, produced from SNP calls in STACKS (b) Distance matrix produced by MinHash distance algorithm in *Mash*. (c) *Reynoutria japonica* (8X) analyzed alone, using Nei’s distance. (d) *Reynoutria japonica* (8X) analyzed alone, using *Mash* distance.

### Extraction and sequencing

We extracted DNA from leaf tissue using a FastPrep homogenizer and a Qiagen Plant Mini kit (Qiagen, Hilden, Germany). We quantified DNA concentration using a Qubit fluorometer and checked quality by running the samples on a 1% agarose gel (Invitrogen; Carlsbad, CA). We sent samples to the Cornell Institute of Biotechnology for library preparation and sequencing. Genotyping-By-Sequencing (GBS) was performed according to the protocol established by Elshire (2011). Briefly, this process fragments DNA using a restriction enzyme (EcoT22I), then ligates barcoded primers to the restriction ends. The barcodes allow samples to be distinguished after the next step, multiplexed PCR. The facility used an Illumina HiSeq to sequence 100 bp single-end reads for all samples.

### Genetic diversity

We used the STACKS (version 2.40) platform to process sequences and call SNPs (Catchen et al. 2011). We first de-multiplexed the reads and removed barcodes using the *process_radtags* script. Since knotweed lacks a reference genome, we used the non-referenced aligned SNP-calling pipeline, *denovo_map*, in STACKS. SNP calling in polyploids is complicated by the presence of several possible alleles at polymorphic loci (reviewed by Blischak et al. 2017, Box 1). To maximize the likelihood of calling accurate SNPs, we used a method similar to that used to call SNPs in tetraploid sturgeon (Anderson et al. 2017). We collected only loci that had a minimum depth of eight (-m 8), a maximum of two mismatches between aligned sequences within an individual (-M 2), and a maximum of one mismatch when comparing to the catalog (-n 1). Since STACKS does not explicitly allow for polyploidy, this method concatenates homeologs. This results in loss of loci due to allele dosage uncertainty, which is the uncertainty over whether copy number is due to sequencing depth or multiple homologous chromosomes (Blischak et al. 2017). To optimize filtering parameters, we ran several iterations of the *populations* module to sequentially test parameters to give the greatest number of polymorphic loci present in 80% of different populations (Paris et al. 2017). We ultimately filtered loci for only SNPs that were present in at least 60% of samples with a coverage above 10x, using a minimum minor allele frequency of 0.05, and a maximum observed heterozygosity of 0.9. This resulted in 12,912 SNPs (sequences publicly available on the NCBI SRA: BioProject PRJNA574173). Genetic diversity estimates of expected heterozygosity (H_e_), nucleotide diversity (π), and percent polymorphic loci were exported from STACKS.

Since genotyping error can cause overestimation of genetic diversity, we checked diversity estimates against a subset of high-confidence SNPs. We re-ran the STACKS SNP calling pipeline with very strict parameters to get a shorter list of 265 SNPs. This subset had 12.7% missing data with average coverage over 60x in all individuals (minimum coverage of 20x at all loci). The high coverage means that it is extremely unlikely that any of these loci represent genotyping errors. We then compared genetic diversity between the entire SNP set to high-confidence loci to account for genotyping error.

### Population structure

We calculated genetic distance between individuals from STACKS-generated SNPs using the *dist*.*genpop* function in *adegenet*, and made the matrix Euclidean using *cailliez* in *ade4* (Jombart 2008; Dray & Dufour 2007). We performed Principal Coordinate Analysis (PCoA) on both distance matrices using the *dudi*.*pco* function in *ade4* (Dray and Dufour 2007). PCoA is often more reliable than Principal Component Analysis (PCA) for GBS data because the latter is more influenced by missing data (Legendre and Legendre 2012). However, to corroborate these patterns, we also performed Principal Component Analysis (PCA) using *pcadapt* (Luu and Blum 2017).

Since SNPs called from polyploid samples may be unreliable, we compared our results to an alternate method for measuring genetic distance known as the MinHash technique. The MinHash technique is a form of probabilistic locality-sensitive hashing (Indyk & Motwani 1998) that reduces the complexity of a dataset to a representative ‘sketch’ (Ondov et al. 2016). In the sketching process, *Mash* converts randomly sampled *k*-mers (sequences of length *k*) from raw sequence reads into computationally efficient ‘hashes’, many of which form a sketch. Sketches can be thought of as genetic summaries and can be compared to find a reliable estimate for the mutation rate between two sequences with relatively low computational load (Ondov et al. 2016). This process takes advantage of the fact that hashes can be more rapidly compared than sequences aligned and, due to random sampling, the degree to which *Mash* can accurately sketch a given sequence scales with the size of the sketch, rather than the size of the genome (Ondov et al. 2016). Though some precision is lost, genetic distances can be estimated without many of the assumptions of other programs that handle genetic data. In particular, *Mash* does not assume diploidy within samples. We sketched each individual using a *k*-mer size of 27 (-k 27) and 1,000,000 hashes (-s 1000000) per individual, omitting *k*-mers that appeared fewer than five times (-c 5) before calculating *Mash* distance. *Mash* produced a distance matrix that we analyzed in parallel to the outputs from STACKS. To test the relationship between the *Mash* distance matrix and our Nei’s distance matrix, we ran a Mantel test using 1,000 permutations.

To visualize phylogenetic relationships, we produced a neighbor-joining (NJ) tree using *phangorn* 2.3.1 (Shliep 2010). We optimized this tree using both maximum parsimony and maximum likelihood (ML) methods through the *parsimony* and *optim*.*pml* functions in *phangorn*. We generated several trees that were compared through log likelihood. We compared models using both AICc and BIC. Bootstrap values were produced by 100 iterations.

We used the program ADMIXTURE 2.40 to detect structure and admixture between populations (Alexander et al. 2009). ADMIXURE, similarly to the commonly used STRUCTURE software (Pritchard et al. 2000), uses multi-locus genotype data to detect populations by assigning individuals to inferred ancestral populations. We ran ADMIXTURE for all values of K between one and thirty-eight (the number of sites sampled) using the default parameters.

### Landscape variables

We tested for evidence of Isolation-by-Distance (IBD) using distance-based redundancy analysis (dbRDA) implemented with the *capscale* function in the *vegan* package (Oksanen et al. 2017) in R. In this analysis, we decomposed the geographic distance matrix into principal components, then tested how the variance in the genetic distance matrix is explained by the spatial eigenvectors using ANOVA (Legendre & Legendre 2012). The dbRDA analysis also allowed testing of different factors to explain variation in the genetic distance matrix, including site, species, sex, and habitat. To determine significance, we used PERMANOVA (Permutational multiple analysis of variance). PERMANOVA is used to compare multivariate groups and tests the null hypothesis that the centroids of the groups are equivalent. Unlike traditional MANOVA, PERMANOVA measures significance by comparing the *F* test result to random permutations of samples from each group (Oksanen et al. 2017). To choose the best fitting model, we used stepwise model selection in the *ordistep* function of *vegan* (Oksanen et al. 2017). The best model was based on a maximum of 50 steps of addition and removal of variables to determine best fit, using permutation *P*-values as an alternative to AIC (Oksanen et al. 2017). In conjunction, we measured the explanatory power of all nineteen Worldclim environmental variables (Table S1) on our two distance matrices using Latent Factor Mixed Models (LFMM) implemented in the *LEA* package in R (Frichot & Francois 2015). LFMMs provide a compromise between power and error rate that is relatively conservative when testing environmental associations (Frichot et al. 2013; Villemereuil et al 2014). Their power lies in the simultaneous testing of environmental correlations while estimating the hidden effect of population structure, so that these are not confounded. The SNPs associated with the environmental variables were determined based on their z-score. Z-score is calculated by the Gibbs sampler algorithm run for 50,000 sweeps after a burn-in period of 10,000 sweeps. The threshold for the z-scores was determined after a Bonferroni correction of α = 0.01. Loci with z-scores > 4.0 and *p* < 10^−5^ were considered significantly associated.

## Results

### Genetic diversity

#### Diversity measures

We found substantial genetic diversity in the populations of *R*. *japonica* we investigated (π = 0.0024, SE = 0.0001; Table 1). Genetic diversity was greater in *R*. *sachalinensis* and *R*. x *bohemica* than in *R*. *japonica* according to heterozygosity measurements (Table 1). Percent polymorphic loci was greatest in the hybrid, and lower for both parent species. Conversely, the largest number of private SNPs were found in the *R*. *sachalinensis* population. The lowest number of private alleles were found in *R*. x *bohemica*.

**Table 1.** STACKS-derived diversity data for individual species. H_e_: Expected Heterozygosity; π: Nucleotide diversity; SE: Standard Error

| Species | Ploidy | # Samples | Total Loci | Polymorphic Loci | % Polymorphic Loci |
| --- | --- | --- | --- | --- | --- |
| Japanese | 8 | 55 | $1.95 \times 10^7$ | 12980 | 0.665 |
| Bohemian | 6 | 18 | $1.10 \times 10^7$ | 11711 | 1.061 |
| Giant | 4 | 15 | $2.05 \times 10^7$ | 21540 | 1.046 |

| | Private SNPs | $H_e$ | SE | $\pi$ variant | $\pi$ invariant |
| --- | --- | --- | --- | --- | --- |
| Japanese | 1091 | 0.279 | 0.002 | $0.283 \pm 0.002$ | $0.0024 \pm 0.0001$ |
| Bohemian | 858 | 0.303 | 0.001 | $0.318 \pm 0.001$ | $0.0035 \pm 0.0001$ |
| Giant | 1617 | 0.309 | 0.001 | $0.326 \pm 0.001$ | $0.0038 \pm 0.0001$ |

We also measured genetic diversity in a subset of high-confidence SNPs. We found that all measures of diversity were comparable in the subset (Table S3). We observed nucleotide diversity of 0.0021 ± 0.0002 for *R*. *japonica*, 0.0023 ± 0.0003 for *R*. x *bohemica*, and 0.0030 ± 0.002 for *R*. *sachalinensis*. While these estimates were lower than what we found for the full SNP set, we found the same pattern, with *R*. *japonica* the least diverse, and *R*. *sachalinensis* the highest.

### Population structure

Using several different approaches, we found little evidence of genetic structure among populations of each of species of knotweed we investigated, although there were genetic differences among the species. We first compared genetic distance matrices produced by two independent methods, SNP-based genotyping and *Mash* distance. We observed a close correlation between the SNP-based Nei’s distance matrix and the *Mash* distance matrix using a Mantel test (1000 permutations, *r* = 0.625, *p* = 0.001). Next, we performed principal coordinates analysis to summarize variation in both matrices. The first and second principal coordinate axes of the PCoA explained 22% and 5.8% of the variation for Nei’s distance and 19% and 3.7%, respectively, for *Mash* distance (Figure 3a,b). The first axis separated samples based largely on species within the hybrid complex. The second and third axes exposed differences in genetic diversity within species for both matrices. However, in the *Mash* distance matrix, the second and third PCs were switched. We saw strong structure based on species and similar clustering (Figure 3a,b) for both matrices. Most sites contained only *R*. *japonica* and therefore, clustered together, revealing very little population-based structure (Figure 3c,d). The sites containing *R*. *sachalinensis* were distinct from *R*. *japonica* samples and from each other. The sites containing *R*. x *bohemica* were the most variable on PC1 and were intermediate between the parental species (Figure 3a,b). All of the observed patterns were consistent between PCoA representations of both distance matrices (Figure 3a,b). The PCoA representation of *R*. *japonica* analyzed alone showed a cluster of samples on the positive side of PC1 that may represent clonal samples (Figure 3c). However, most sites separated on both PCs and sites did not form distinct groups.

**Figure 3.**
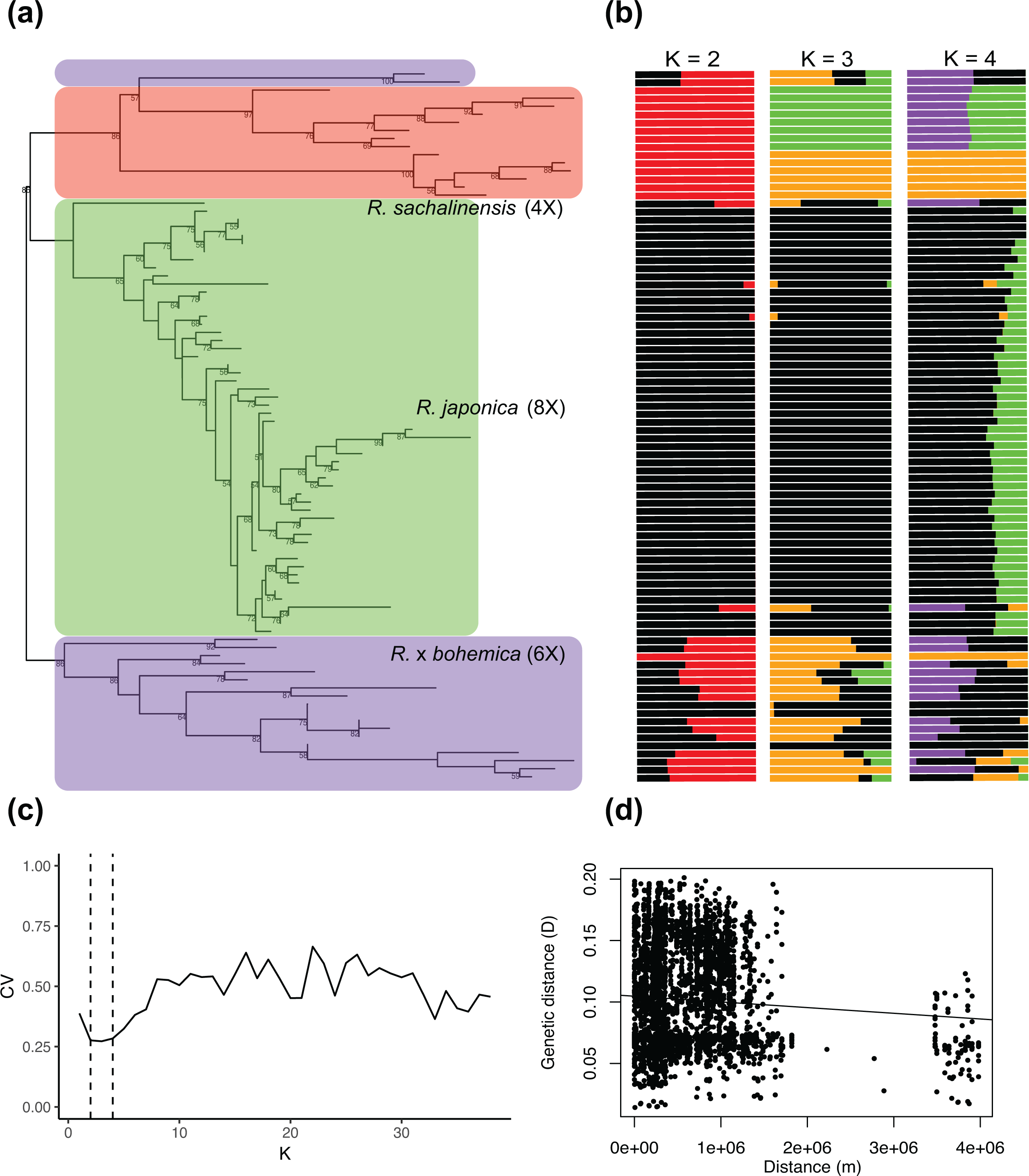
Maximum likelihood tree and ADMIXTURE results. (a): ML tree from SNP calls. *R*. *sachalinensis* is shown in red, *R*. *japonica* in green, and *R*. x *bohemica* in blue. Bootstrap values higher than 50 are shown at nodes. (b): Admixture assuming K = 2, 3, and 4 ancestral populations. Each bar represents one sample. (c) Coefficient of variation (CV) for each tested value of K for admixture analysis. Dashed lines show K = 2-4. (d) Geographic and Nei’s genetic distance (D) correlation.

Our ML trees mostly corroborated the results we found in the principal coordinate analysis. In particular, *R*. *japonica* populations were not differentiated, but both *R*. *sachalinensis* and *R*. x *bohemica* populations showed significant site-based differentiation (Figure 4a).

Bootstrap values were low for most branches within the *R*. *japonica* clade, but high within the other clades. Notably, the tree highlights a strong isolation of two particular samples at the top of the phylogeny, which were male *R*. x *bohemica* within a site containing mostly female *R*. *japonica*.

We examined ADMIXTURE values of K from 1-38 and found the best support for K = 2 (Figure 4b, c). Consistent with other analyses, we saw distinction of *R*. *japonica* and *R*. *sachalinensis* and admixture in sites that contained *R*. x *bohemica* hybrids. This is most apparent when K = 2. The Cornell population is grouped with *R*. *sachalinensis* at K = 2, but shown to be more similar to the *R*. x *bohemica* group when K is increased to 3. The homogenous block of *R*. *japonica* was not separated for any level of K less than 10. At levels above 10, individuals were assigned to differing inferred ancestral populations, but these did not correspond to collection site. Analysis of *R*. *japonica* in isolation did not return different patterns than when analyzed in conjunction with other species.

### Landscape variables

We did not find evidence for Isolation-by-Distance (IBD) because there was no relationship between geographic distance and genetic distance, as shown by dbRDA (*F* = 0.907, *p* = 0.679; Table 2, Figure 4d). This pattern was not affected by the distance measure used, as there was similarly low explanation of the variation in the *Mash* genetic distance by geographic distance (*F* = 1.03, *p* = 0.387; Table 2). However, the combination of factors that best explained genetic variation in PERMANOVA differed between measures of genetic distance. While both models included species and site, Nei’s distance was partially explained by habitat, and *Mash* distance by sex, though the marginal effects of each of these factors were weak relative to species and site (Table 2).

**Table 2.** PERMANOVA results for two genetic distance estimation methods, SNP-based Nei’s distance, and *Mash* distance from the *Mash* MinHash algorithm. Permutational model selection determined the best model to explain the Nei’s distance matrix as Species + Site + Habitat, and the *Mash* distance matrix as Species + Site + Sex. *F*- and *p*-values shown are marginal values when each factor is tested against all other factors. Bold values were significant at α = 0.05, italicized values are marginally significant at α = 0.1

| Nei distance |  |  | *Mash* distance |  |  |
| --- | --- | --- | --- | --- | --- |
| Factor | *F* | *p* | Factor | *F* | *p* |
| Species | **7.31** | **0.001** | Species | **2.83** | **0.001** |
| Site | **1.56** | **0.009** | Site | **1.19** | **0.001** |
| *Habitat* | *2.29* | *0.064* | *Sex* | *1.30* | *0.071* |
| Overall | **3.38** | **0.001** | Overall | **1.87** | **0.001** |
| Sex | 0.729 | 0.715 | Habitat | 1.08 | 0.263 |
| Distance | 0.907 | 0.679 | Distance | 1.03 | 0.387 |

We examined correlations with environmental variables using Latent Factor Mixed Models. However, loci were not significantly correlated with any of nineteen climatic variables (after correction for multiple testing; all values of *p* > 0.9).

## Discussion

### Overview

In this study, we examined one of the worlds’ most successful and widespread weeds using next-generation sequencing and genomic analyses. In contrast to the expectation that invasive species in general should have low genetic diversity because of founder effects, and also contrary to prior studies indicating the near absence of genetic diversity in this group, we found substantial genetic diversity in the Japanese knotweed species complex in the North American populations we sampled. However, there was little population structure, and the distribution of genetic diversity was not correlated with climatic or geographic factors in *R*. *japonica*. Our study highlights the importance of careful assessment of genetic diversity and consideration of the role multiple introductions and hybridization can take in species invasions. This work also provides an illustration of how contemporary analytical techniques can be employed to evaluate population structure of polyploid species lacking a reference genome.

### Differences in genetic diversity between hybrid species

Due perhaps to its ubiquity and invasiveness, *R*. *japonica* has been the focus of several genetic studies over the last two decades in North America. While sexual reproduction was known to occur in *R*. *japonica* populations, most studies assumed that the majority of reproduction occurred through asexual means (Forman & Kesseli 2003; Richards et al. 2012).

Rather than completely a simple binary clonal versus non-clonal designation, researchers have recognized that the genetic definition of clonality is a continuum of similarity (Bailleul et al. 2016). Strategies for dividing samples into clonal groups, or multi-locus lineages, usually involve designating a minimum threshold for genetic distance that corresponds to an estimate of sequencing error and somatic mutation (Kamvar et al. 2015). In the absence of prior knowledge of this threshold, this strategy relies on a frequency distribution of genetic distances that often shows a peak near zero designating clonal lineages (Meirmans & van Tienderen 2004). We can reject the hypothesis that North American *R*. *japonica* are entirely clonal, since even with a conservative threshold for minimum clonal distance we would observe several clonal linages; most samples showed moderate genetic distance (D; range 0.014-0.201; median: 0.094). Studies using SNP markers have observed similar genetic distances of between 0.02 and 0.19 for subpopulations of sexual *Solanum lycopersicum* (Sim et al. 2015). Further, the level of genetic diversity we measured (π = 0.0024) is comparable to what has been seen in other polyploid species (Cornille et al. 2016), and much higher than what has been measured in clones (Gutekunst et al. 2018). Because both genetic distances and overall diversity were higher than would be expected for purely clonal lineages, we suggest that stands of *R*. *japonica* often contain related individuals in addition to clones.

Precise estimation of genetic diversity is problematic in polyploid organisms because estimated heterozygosity can be inflated due to homologous chromosomes yielding homeologous sequences (Blischak et al. 2017). In addition, taxa that undergo whole genome duplication often then undergo diploidization, a process by which homeologs are lost and diploid inheritance is resumed (Barker et al. 2016). If the SNPs are mostly clustered in non-coding regions or on silenced homeologous chromosomes, different genotypes may be functionally identical (Mandáková et al. 2017). Further, somatic mutations occur often in plants, and can play an important role in evolution, especially in clones (Whitham & Slobodchikoff 1981; Schoen & Schultz 2019). While it is conceivable that some of the genetic diversity we observe is due to somatic mutations within clones, it is unlikely to be a major source of genetic diversity in introduced areas, given the relatively short time since *Reynoutria spp*. were introduced to North America. Even if somatic mutations did contribute to genetic diversity in this complex, this would not contradict our finding of substantial genetic diversity, but would provide one mechanism for the generation of this diversity.

We took several measures to ensure that genetic diversity estimates we report are not due to artifacts of sequencing or data analysis, including conducting the analyses with a subset of high-confidence SNPs and confirming the results with *Mash* analysis. The congruence of the analysis on the high-confidence SNP data and the full dataset indicates that the patterns we found are unlikely to be due to artifacts or error. While the values we report for π and H_s_ are not necessarily comparable to those for a diploid species, they represent a useful estimation of relative genetic diversity. *Mash* estimation is less influenced by ploidy than alignment-based SNPs, so it is a useful benchmark to show that within-population diversity is substantial. Thus, it is unlikely that polyploidy prevented this study from correctly identifying the pattern of genetic diversity.

There is a somewhat surprising pattern in *Reynoutria* species of higher-ploidy species exhibiting lower genetic diversity (Hollingsworth et al. 2000; Richards et al. 2012). Even if they are not entirely clonal, octoploid *R*. *japonica* show less nucleotide diversity than either tetraploid *R*. *sachalinensis* or hexaploid *R*. x *bohemica*. Polyploid individuals have inherently higher allelic diversity than diploids, since a duplicated genome allows twice the potentially expressed alleles.

However, at the population level polyploid populations often have lower genetic diversity owing to life history constraints. In a striking recent example, a triploid genotype of marbled crayfish (*Procambarus virginalus*) have been successful in invading ecosystems worldwide (Gutekunst et al. 2018), despite complete genetic identity. The pattern is also evident in native species colonizing new areas. Native Pacific Northwest hawthorns (*Crataegus spp*.) show high genetic diversity throughout a species complex, but several clonal polyploid lineages have colonized wider ranges than their diploid ancestors (Coughlan et al. 2017). Similarly, octoploid *R*. *japonica* shows both higher asexual reproduction and wider range than tetraploid *R*. *sachalinensis*. There is a well-established association between polyploidy and asexual reproduction in both plants and animals (Otto & Whitton 2000; Neiman et al. 2014). However, the reasons for the repeated pattern of range expansion following asexual polyploidization are not completely understood.

One possible explanation is that polyploidization allows a species access to a new niche (Baniaga et al. 2020), and asexual reproduction allows the neopolyploid a means to reproduce in the absence of mates. The polyploid history of *R*. *japonica* has not yet been studied, so a relationship between niche divergence and polyploidization cannot be determined. Future studies may be able to exploit this polyploid series to better understand the importance of ploidy to plant colonization.

### Population Structure

Population structuring, like diversity, differed between *Reynoutria* species. The difference between species was unsurprisingly the strongest signal of differentiation, but there was also some structuring by collection site for *R*. *sachalinensis* and *R*. x *bohemica*. Within *R*. *japonica*, we saw no evidence of population structuring (Figure 3). While this could be evidence of panmixia, wherein gene flow is high between all sites, the wide geographic range of sites means that this explanation is highly unlikely (Waples & Gaggiotti 2006). Instead, we hypothesize that the *R*. *japonica* samples we analyzed were derived from a homogenous source population, and have not yet genetically differentiated within the invaded range. This interpretation is corroborated by undetectable Isolation-by-Distance (IBD) in these populations, as well as the stochastic nature of *R*. *japonica* introductions to North America (Barney 2006). Within the hybrid and *R*. *sachalinensis*, there was strong structuring, as was seen in other studies (Grimsby et al.

2009; Richards et al. 2012; Gaskin et al. 2014). Depending on the analysis used, we saw slightly differing patterns. While PCoA clearly distinguishes Cornell and Maine from all other sites, the ML tree included several *R*. x *bohemica* from CT as closely related to Cornell and Maine (two samples at the top of the tree). One possible explanation is that these CT samples are introgressed backcrosses between *R*. x *bohemica* and *R*. *sachalinensis*, as has been found previously in American populations (Gammon et al. 2010).

Another notable pattern in population structure is the confounding factor of sex. *Reynoutria* are subdioecious, with females and hermaphrodite individuals that rarely produce seeds (*i*.*e*. function as males; Forman & Kesseli 2003). The clonal UK population of *R*. *japonica* consists entirely of females, though male *R*. *sachalinensis* and *Fallopia balduschiana* often provide pollen for hybridization (Hollingsworth & Bailey 2000; Bailey 2013). In contrast, male *R*. *japonica* are present (if rare) in North America (Forman & Kesseli 2003). Male *R*. *sachalinensis* and *R*. *bohemica* are far more common, and account for the entire Maine population as well as two samples which show great divergence from the CT population (Figure 4a,b). Biased sex ratios impact N_e_ and therefore may have wide-ranging effects on estimation of genetic diversity. Biased sex ratios should be selected against, since the rarer sex has a comparative fitness advantage (Fisher 1958). Biased ratios can persist due to several factors, including parthenogenesis as in several vertebrates (Lynch 1984), *Wolbachia* infection in insects (Stouthamer et al. 1999), and partial reproductive isolation variation in plants (Barnard-Kubow & Galloway 2017), to name a few. In the case of knotweed, it may simply be the case that the rhizomes that were moved to Europe and North America were more commonly female than male, and that populations have not have had time to equalize.

### Landscape variation

Unlike several other population genomic studies in invasive species (Cornille et al. 2016; Trumbo et al. 2016; Combs et al. 2018), we did not find evidence of Isolation-By-Distance (IBD). While the introduction history of *Reynoutria spp*. is complicated, lack of IBD for a widespread species with no clear dispersal boundaries is fairly surprising. PERMANOVA showed collection site-based structure, which indicates that there is some geographic component to genetic structure. However, the fact that the populations do not show IBD indicates that these populations are scattered across the landscape, likely due to a stochastic introduction history rather than linear spread. In addition, we also observed no correlations between genetic variation and climatic variables. There are several possible explanations for this finding. The LFMM method uses a conservative correction for multiple testing that may result in values below the significance limit (Frichot et al. 2013). However, PERMANOVA indicates that *R*. *japonica* has been introduced stochastically across the landscape, and this may result in little geographic differentiation. While we found evidence of sexual reproduction, frequent clonal propagation of *Reynoutria spp*. likely plays a strong role in both preventing IBD and minimizing environmental correlations (Reusch et al. 2000). Research on clonal invasive cogongrass (*Imperata cylindrica*) showed similar lack of IBD, presumably due to limitations in diversity (Burrell et al. 2015). In the *R*. *japonica* native range, there is predictable species separation, but no information on hybrids. *R*. *sachalinensis* grows mostly in the northern Sakhalin island in Russia and Hokkaido, the northernmost island in Japan (Inamura 2000). Although the PERMANOVA did not detect an overall effect of distance, we only found *R*. *sachalinensis* in the North (Figure 1), suggesting a pattern that might be detected with wider sampling. The lack of landscape-scale patterns may be a product of both life history and invasion history. In outcrossing populations, adaptation along a linear gradient promotes differentiation (Rousset 1997, Vekemans & Hardy 2004), but populations that show clonal reproduction diverge genetically much more slowly during spread. As a compounding factor, multiple introductions of *Reynoutria spp*. (Barney 2006) have likely led to a complicated mosaic of genetic diversity across the landscape that is challenging to untangle at this level of resolution.

### Population genetics of polyploids

There are several distinct challenges of inferring population genetic patterns from polyploid data. In general, sequencing to sufficient depth to capture allele variation is expensive, and large genome size for many polyploids, including *R*. *japonica*, further complicates matters. A locus that is partially heterozygous creates uncertainty in dosage, which complicates accurate inference of allele frequencies (Meirmans et al. 2018). In addition, most population genetic theory is built around outcrossing diploid models that may or may not apply to clonal polyploids, including simple measurements such as fixation index (F_ST_; Dufresne et al. 2014; Meirmans et al. 2018). At each stage of analysis (genotyping, estimation of population structure, and landscape associations), we used parallel analysis techniques to account for some of this uncertainty, and relied on relative measures of genetic diversity whenever possible to avoid bias from genotyping uncertainty. Across analyses, we found broadly similar results, supporting our final conclusions.

Whole-genome sequencing is becoming economically feasible for a wider array of polyploid species, so may soon replace reduced-representation methods (e.g. Edger et al 2019). However, reduced-representation methods remain an excellent means of capturing variation in many markers for large sample sizes. To understand coarse patterns in data, *Mash* and similar k-mer hashing techniques may be particularly useful due to the few assumptions required and the low computational load (Ondov et al. 2016; Viochek & Weigel 2019). However, it lacks many of the data-cleaning steps of other genotyping platforms and therefore may be prone to overestimating genetic diversity. The *STACKS* platform has numerous well-curated tools for many applications. For projects with many markers, however, a high-performance computer is needed. Population genetic studies in polyploid species will always face challenges, but bioinformatic tools are increasingly accommodating higher ploidy levels.

### Conclusions

Japanese knotweed populations in North America display a complex genetic structure that differs from what has been seen at smaller scales with other genetic tools. Unlike what has been traditionally understood for North America, and what has been seen in Great Britain, there is substantial genetic diversity within populations of knotweed. While clonal spread is undoubtedly important to the invasive success of knotweed, it is evident that sexual reproduction has also occurred during its rapid spread due to the genetic variation that exists with sites, although somatic mutation could have potentially also played some role in creating diversity. The lack of evidence for IBD in knotweed populations in the present study is likely linked to its stochastic introduction throughout the late 19th and early 20th centuries, and its frequent clonal propagation and less frequent sexual reproduction. We provide evidence that knotweed has avoided the genetic paradox of invasions (rapid spread despite loss of genetic variation) by two means. First, populations that were assumed to be clonal harbor levels of heterozygosity that indicate there has not been complete loss of diversity in introduced populations, either because of sexual reproduction or retained polyploid diversity. While these populations have not differentiated across the landscape, phenotypic variability may not be limited by lack of genetic variation (Richards et al. 2012). Second, hybridization with *R*. *sachalinensis* occurs readily, so the species complex as a whole can be said to avoid the genetic paradox through hybridization. Since polyploidy can play a role in both harboring heterozygosity and allowing hybridization, we expect that it has influenced knotweed invasion. Finally, we have shown that GBS data can be viable for non-model polyploids by comparing our results between two fundamentally different methods for measuring genetic differentiation. Taken together, our results provide insights into the roles of factors such as clonality and hybridization in shaping the population genomics and success of an invasive species, and more broadly help to inform our understanding of the process of colonization.

## Acknowledgements

We thank D. Fogarasi, S. Davey, G. Diaz, and J. Goehl for field assistance; A. VanWallendael, P. VanWallendael, K. Ansaldi, J. Huffstetler, B. Wade, Dr. N. Bauer, Dr. J. Weber, and Dr. N. Muth for sending samples; D. Fogarasi and J. Park for lab assistance; M. Combs and E. Puckett for assistance with data analyses; and Dr. Jason Munshi-South, Dr. John Wehr, Dr. James Lewis, Dr. Beth Ansaldi, Dr. Michael Sekor for helpful comments on the manuscript. We acknowledge the following field stations and reserves: Louis Calder Center, Goodwin College, USFS, Hubbard Brook Experimental Forest, Mohonk Reserve, Blandy Experimental Farm, Wells Reserve, Cornell University, New York Botanical Garden, Claytor Nature Study Center. We thank the Fordham Graduate School of Arts and Sciences, Fordham Graduate Student Association, and Louis Calder Center for funding.

## Author Contributions

AV designed the study, collected and analyzed data and wrote the manuscript. MA analyzed data and wrote the manuscript. SJF contributed to study design and writing of the paper.

## Data Availability

Sequences have been deposited into the NCBI Short Read Archive (SRA) as BioProject PRJNA574173

