## Supplemental data for "Patterns of population genomic diversity in the invasive Japanese knotweed species complex"

**Supplemental Material**

**Table S1**. Bioclim variables used in LFMM (Latent Factor Mixed Models)

| Name | Variable |
| --- | --- |
| BIO1 | Annual Mean Temperature |
| BIO2 | Mean Diurnal Range (Mean of monthly (max temp - min temp)) |
| BIO3 | Isothermality (BIO2/BIO7) (* 100) |
| BIO4 | Temperature Seasonality (standard deviation *100) |
| BIO5 | Max Temperature of Warmest Month |
| BIO6 | Min Temperature of Coldest Month |
| BIO7 | Temperature Annual Range (BIO5-BIO6) |
| BIO8 | Mean Temperature of Wettest Quarter |
| BIO9 | Mean Temperature of Driest Quarter |
| BIO10 | Mean Temperature of Warmest Quarter |
| BIO11 | Mean Temperature of Coldest Quarter |
| BIO12 | Annual Precipitation |
| BIO13 | Precipitation of Wettest Month |
| BIO14 | Precipitation of Driest Month |
| BIO15 | Precipitation Seasonality (Coefficient of Variation) |
| BIO16 | Precipitation of Wettest Quarter |
| BIO17 | Precipitation of Driest Quarter |
| BIO18 | Precipitation of Warmest Quarter |
| BIO19 | Precipitation of Coldest Quarter |

**Table S2**: Sample information

| Sample | Site | Subsite | Latitude | Longitude | Species | Sex | Habitat |
| --- | --- | --- | --- | --- | --- | --- | --- |
| BM4 | Adirondack | BM1 | 43.87477 | -74.43083 | japonica | unknown | edge |
| BM6 | Adirondack | BM2 | 43.84893 | -74.41985 | japonica | unknown | edge |
| HU2 | Adirondack | HU | 43.66835 | -73.96103 | japonica | unknown | road |
| IL1 | Adirondack | IL | 43.78333 | -74.24922 | japonica | unknown | edge |
| IN1 | Adirondack | IN | 43.76536 | -74.84400 | japonica | unknown | edge |
| LG1 | Adirondack | LG1 | 43.45692 | -73.69495 | japonica | unknown | road |
| LG2 | Adirondack | LG2 | 43.47268 | -73.68735 | japonica | unknown | road |
| RA1 | Adirondack | RA | 43.74420 | -74.89607 | japonica | unknown | edge |
| TL1 | Adirondack | TL | 43.52803 | -73.70226 | japonica | unknown | road |
| TL2 | Adirondack | TL | 43.52781 | -73.70352 | japonica | unknown | road |
| 102 | Cornell | Cornell1 | 42.46592 | -76.43087 | sachalinensis | unknown | river |
| 103 | Cornell | Cornell1 | 42.46592 | -76.43087 | sachalinensis | unknown | river |
| 104 | Cornell | Cornell1 | 42.46592 | -76.43087 | sachalinensis | unknown | river |
| 108 | Cornell | Cornell2 | 42.45989 | -76.43189 | sachalinensis | unknown | edge |
| 110 | Cornell | Cornell2 | 42.45989 | -76.43189 | sachalinensis | unknown | edge |
| 121 | Cornell | Cornell3 | 42.46835 | -76.41454 | sachalinensis | unknown | edge |
| 125 | Cornell | Cornell3 | 42.46835 | -76.41454 | sachalinensis | unknown | edge |
| NYBG | CT | Bronx | 40.85693 | -73.87717 | japonica | unknown | river |
| 74 | CT | Calder | 41.12851 | -73.73053 | bohemica | m | edge |
| 75 | CT | Calder | 41.12851 | -73.73053 | bohemica | m | edge |
| 67 | CT | Calder | 41.12866 | -73.73060 | japonica | f | edge |
| 70 | CT | Calder | 41.12866 | -73.73060 | japonica | f | edge |
| Seed_A | CT | Calder | 41.12747 | -73.73091 | japonica | seed | edge |
| Seed_B | CT | Calder | 41.12747 | -73.73091 | japonica | seed | edge |
| 91 | CT | CT | 41.06407 | -73.54607 | bohemica | unknown | river |
| 94 | CT | CT | 41.06407 | -73.54607 | bohemica | unknown | river |
| 95 | CT | CT | 41.06407 | -73.54607 | bohemica | unknown | river |
| 82 | CT | Goodwin1 | 41.73460 | -72.63804 | bohemica | unknown | edge |
| 83 | CT | Goodwin1 | 41.73460 | -72.63804 | bohemica | unknown | edge |
| 77 | CT | Goodwin1 | 41.73452 | -72.64038 | japonica | unknown | river |
| 79 | CT | Goodwin1 | 41.73452 | -72.64038 | japonica | unknown | river |
| 87 | CT | Goodwin2 | 41.74587 | -72.64262 | bohemica | unknown | river |
| 88 | CT | Goodwin2 | 41.74587 | -72.64262 | bohemica | unknown | river |
| 90 | CT | Goodwin2 | 41.74587 | -72.64262 | bohemica | unknown | river |
| 38 | ME | Wells1 | 43.34492 | -70.55291 | sachalinensis | m | river |
| 39 | ME | Wells1 | 43.34492 | -70.55291 | sachalinensis | m | river |
| 42 | ME | Wells2 | 43.34686 | -70.54937 | sachalinensis | m | river |
| 43 | ME | Wells2 | 43.34686 | -70.54937 | sachalinensis | m | river |
| 44 | ME | Wells2 | 43.34686 | -70.54937 | sachalinensis | m | river |
| 48 | ME | Wells3 | 43.34481 | -70.56062 | sachalinensis | m | road |
| 49 | ME | Wells3 | 43.34481 | -70.56062 | sachalinensis | m | road |
| 50 | ME | Wells3 | 43.34481 | -70.56062 | sachalinensis | m | road |
| 51 | Mid Atlantic | Blandy | 39.08063 | -78.08574 | japonica | f | road |
| 54 | Mid Atlantic | Blandy | 39.08063 | -78.08574 | japonica | f | road |
| 55 | Mid Atlantic | Blandy | 39.08063 | -78.08574 | japonica | f | road |
| 156 | Mid Atlantic | Claytor | 37.38097 | -79.54347 | japonica | f | river |
| 158 | Mid Atlantic | Claytor | 37.38097 | -79.54347 | japonica | f | river |
| 160 | Mid Atlantic | Claytor | 37.38097 | -79.54347 | japonica | f | river |
| 18 | Mid Atlantic | Delaware | 40.60519 | -75.19276 | japonica | f | river |
| 19 | Mid Atlantic | Delaware | 40.60519 | -75.19276 | japonica | f | river |
| 20 | Mid Atlantic | Delaware | 40.60519 | -75.19276 | japonica | m | river |
| 23 | Mid Atlantic | Mohonk | 41.73409 | -74.18843 | japonica | f | edge |
| 24 | Mid Atlantic | Mohonk | 41.73409 | -74.18843 | japonica | f | edge |
| 25 | Mid Atlantic | Mohonk | 41.73409 | -74.18843 | japonica | f | edge |
| 56 | Mid Atlantic | NJ1 | 39.95110 | -74.96804 | japonica | f | lake |
| 57 | Mid Atlantic | NJ1 | 39.95110 | -74.96804 | japonica | f | lake |
| 58 | Mid Atlantic | NJ1 | 39.95110 | -74.96804 | japonica | f | lake |
| 62 | Mid Atlantic | NJ2 | 39.95020 | -74.95555 | japonica | f | lake |
| 64 | Mid Atlantic | NJ2 | 39.95020 | -74.95555 | japonica | f | lake |
| 65 | Mid Atlantic | NJ2 | 39.95020 | -74.95555 | japonica | f | lake |
| CAR | Midwest | Carbondale | 37.71372 | -89.21968 | japonica | unknown | road |
| CHI | Midwest | Chicago | 41.91863 | -87.67090 | japonica | unknown | road |
| MN | Midwest | MN | 44.97108 | -93.26441 | japonica | f | road |
| 127 | NC | NC1 | 36.07876 | -82.34724 | bohemica | unknown | river |
| 132 | NC | NC1 | 36.07876 | -82.34724 | bohemica | unknown | river |
| 133 | NC | NC1 | 36.07876 | -82.34724 | bohemica | unknown | river |
| 139 | NC | NC2 | 35.88135 | -82.77004 | bohemica | unknown | river |
| 144 | NC | NC2 | 35.88135 | -82.77004 | japonica | unknown | river |
| 145 | NC | NC2 | 35.88135 | -82.77004 | japonica | unknown | river |
| 147 | NC | NC3 | 35.83878 | -82.75613 | bohemica | unknown | river |
| 152 | NC | NC3 | 35.83878 | -82.75613 | bohemica | unknown | river |
| 153 | NC | NC3 | 35.83878 | -82.75613 | bohemica | unknown | river |
| 26 | NH | Hubbard1 | 43.93716 | -71.68359 | japonica | f | edge |
| 27 | NH | Hubbard1 | 43.93716 | -71.68359 | japonica | f | edge |
| 30 | NH | Hubbard1 | 43.93716 | -71.68359 | japonica | f | edge |
| 31 | NH | Hubbard2 | 43.66835 | -73.96103 | japonica | f | edge |
| 34 | NH | Hubbard2 | 43.66835 | -73.96103 | japonica | f | edge |
| 35 | NH | Hubbard2 | 43.66835 | -73.96103 | japonica | f | edge |
| WA | West | WA | 47.70664 | -122.08922 | bohemica | unknown | road |
| 1 | WV | WV1 | 37.95254 | -81.07512 | japonica | f | road |
| 3 | WV | WV1 | 37.95254 | -81.07512 | japonica | f | road |
| 5 | WV | WV1 | 37.95254 | -81.07512 | japonica | f | road |
| 10 | WV | WV2 | 37.92733 | -81.11951 | japonica | f | road |
| 6 | WV | WV2 | 37.92733 | -81.11951 | japonica | f | road |
| 8 | WV | WV2 | 37.92733 | -81.11951 | japonica | f | road |
| 12 | WV | WV3 | 37.80961 | -80.92438 | japonica | f | river |
| 13 | WV | WV3 | 37.80961 | -80.92438 | japonica | f | river |
| 15 | WV | WV3 | 37.80961 | -80.92438 | japonica | f | river |

**Table S3:** Genetic diversity estimates using a subset of 265 high-confidence SNPs

| Species | Ploidy | # Samples | Total loci | Polymorphic Loci | % Polymorphic Loci |
| --- | --- | --- | --- | --- | --- |
| Japanese | 8 | 55 | 62447 | 340 | 0.544 |
| Bohemian | 6 | 18 | 39709 | 209 | 0.526 |
| Giant | 4 | 15 | 126551 | 798 | 0.631 |
| Species | Private SNPs | H_e_ | SE | **π** _variant_ | **π** _invariant_ |
| Japanese | 7 | 0.0021 | 0.0001 | 0.353±0.07 | 0.0021±0.0003 |
| Bohemian | 20 | 0.0022 | 0.0002 | 0.419±0.16 | 0.0023±0.0002 |
| Giant | 23 | 0.0029 | 0.0001 | 0.461±0.04 | 0.0030±0.0002 |
